# Ascorbate deficiency does not limit non-photochemical quenching in *Chlamydomonas reinhardtii*

**DOI:** 10.1101/813766

**Authors:** André Vidal-Meireles, Dávid Tóth, László Kovács, Juliane Neupert, Szilvia Z. Tóth

## Abstract

Ascorbate (vitamin C) plays essential roles in development, signaling, hormone biosynthesis, regulation of gene expression, stress resistance and photoprotection. In vascular plants, violaxanthin de-epoxidase (VDE) requires ascorbate (Asc) as reductant, thereby it is required for the energy-dependent component of non-photochemical quenching (NPQ). In order to assess the role of Asc in NPQ in green algae, which are known to contain low amounts of Asc, we searched for an insertional *Chlamydomonas reinhardtii* mutant affected in the *VTC2* gene, essential for Asc biosynthesis. The *Crvtc2-1* knockout mutant was viable and, depending on the growth conditions, it contained 10 to 20% Asc relative to its wild type. When Chlamydomonas was grown photomixotrophically at moderate light, the zeaxanthin-dependent component of NPQ emerged upon strong red illumination both in the *Crvtc2-1* mutant and in its wild type. Deepoxidation was unaffected by Asc deficiency, demonstrating that the Chlorophycean VDE found in Chlamydomonas does not require Asc as a reductant. The rapidly induced, energy-dependent NPQ component, characteristic of photoautotrophic Chlamydomonas cultures grown at high light, was not limited by Asc deficiency either. On the other hand, a reactive oxygen species-induced photoinhibitory NPQ component was greatly enhanced upon Asc deficiency, both under photomixotrophic and photoautotrophic conditions. These results demonstrate that Asc has distinct roles in NPQ formation in Chlamydomonas than in vascular plants.

**One-sentence summary:** In Chlamydomonas -in contrast to seed plants-, ascorbate is not required for violaxanthin deepoxidation and energy-dependent non-photochemical quenching but it mitigates photoinhibitory quenching.

## Introduction

Ascorbate is a multifunctional metabolite, essential for a range of cellular processes in green plants, including cell division, stomatal movement, the synthesis of various plant hormones, epigenetic regulation and reactive oxygen species (ROS) scavenging (Asada, 2006; Foyer and Shigeoka, 2011; Smirnoff, 2018). Within the chloroplast, Asc may also act as an alternative electron donor to photosystem II (PSII) and to PSI (Ivanov et al., 2007; Tóth et al., 2009; Tóth et al., 2011). In vascular plants, violaxanthin de-epoxidase (VDE) requires ascorbate as reductant, thereby Asc plays an essential role in the process of non-photochemical quenching (NPQ) to dissipate excess energy as heat (Bratt et al., 1995; Saga et al., 2010; Hallin et al., 2016).

In order to fulfill the multiple physiological roles of Asc (reviewed by Tóth et al., 2018; Smirnoff, 2018), vascular plants maintain their Asc concentration at a high, approximately 20 to 30 mM level (Zechmann et al., 2011), which is also relatively constant, usually with no more than two-fold increase upon stress treatments and moderate decrease during dark periods (Dowdle et al., 2007). Notwithstanding, Asc concentration may be limiting under environmental stress conditions, as shown by an increased oxidative stress tolerance of plants overexpressing dehydroascorbate reductase, playing an essential role in Asc regeneration (Wang et al., 2010). Regarding NPQ, it was shown that Asc-deficient Arabidopsis plants have slowly inducible and diminished NPQ, whereas Asc-overproducing plants possess enhanced NPQ relative to wild type plants, meaning that Asc may limit the conversion of violaxanthin to zeaxanthin *in vivo* (Müller-Moulé et al., 2002; Tóth et al., 2011). Ascorbate deficient plants are also sensitive to high light, especially in combination with zeaxanthin deficiency (Müller-Moulé et al., 2003).

Green algae, for instance *Chlamydomonas reinhardtii*, produce Asc in a very small amount under favorable environmental conditions (approx. 100 to 400 µM, Gest et al., 2013), and boost it only in case of need, for instance upon a sudden increase in light intensity and in nutrient deprivation (Vidal-Meireles et al., 2017; Nagy et al., 2018). The mode of regulation of Asc biosynthesis differs largely between plants and Chlamydomonas: in contrast to vascular plants, i) green algal Asc biosynthesis is directly regulated by ROS, ii) it is not under circadian clock control and, iii) instead of a negative feedback regulation, there is a feedforward mechanism on the expression of the key Asc biosynthesis gene, *VTC2* (*Cre13.g588150*) by Asc in the physiological concentration range (Vidal-Meireles et al., 2017).

Regarding NPQ, it was described that the violaxanthin deepoxidase found in Chlorophyceae (CVDE) is not homologous to plant VDE but related to a lycopene cyclase of photosynthetic bacteria (Li et al., 2016a). Chlamydomonas CVDE (CrCVDE), encoded by *Cre04.g221550*, has a FAD-binding domain and it is located on the stromal side and not in the thylakoid lumen, as it is the case for plant-type VDE (Li et al., 2016a). The cofactor or reductant requirement of the CrCVDE enzyme has not been investigated, and it is not known whether its activity requires Asc, either directly or indirectly.

Due to the major differences in Asc contents, the regulation of Asc biosynthesis and the VDE enzymes of vascular plants and Chlorophyceae, we decided to assess the role of Asc in the various NPQ components in *Chlamydomonas reinhardtii*. To this end, we characterized an insertional *VTC2* mutant procured from the CLiP library (Li et al., 2016b), possessing only 10 to 20% Asc relative to its parent strain. We have found that, in contrast to vascular plants, Asc deficiency does not limit energy-dependent quenching (qE) and violaxanthin de-epoxidation in Chlamydomonas; instead, Asc deficiency leads to enhanced photoinhibitory quenching (qI) upon excessive illumination.

## Results

### Identification and initial characterization of an Asc-deficient VTC2 insertional mutant of C. reinhardtii and its genetic complementation

To investigate the function of Asc in NPQ in *C. reinhardtii*, we searched for insertion mutants for the *VTC2* gene in the CLiP library (Li et al., 2016b). We found one putative *VTC2* mutant (strain *LMJ.RY0402.058624*, hereafter called *Crvtc2-1*), holding one insertion of the paromomycin resistance (CIB1) casette at the junction site of exon 3 and the adjacent upstream intron of the *VTC2* gene (Fig. 1A). The other available mutants were affected in the 3’UTR region of the *VTC2* gene and/or had multiple insertions in genes other than *VTC2*. One of the strains carrying a CIB1 casette in the 3’UTR region of the *VTC2* gene with an insertion confidence of 95% (*LMJ.RY0402.146784*) was tested, but we found that it had wild-type level Asc content (data not shown); therefore, it was not used for further analyses. Due to the lack of another, independent CIB insertional mutant line affecting only the *VTC2* gene, we carried out several NPQ measurements on our previously published *VTC2*-amiRNA line (Vidal-Meireles et al., 2017) to confirm our findings on the consequences of Asc deficiency on NPQ (see below).

**Figure 1.**
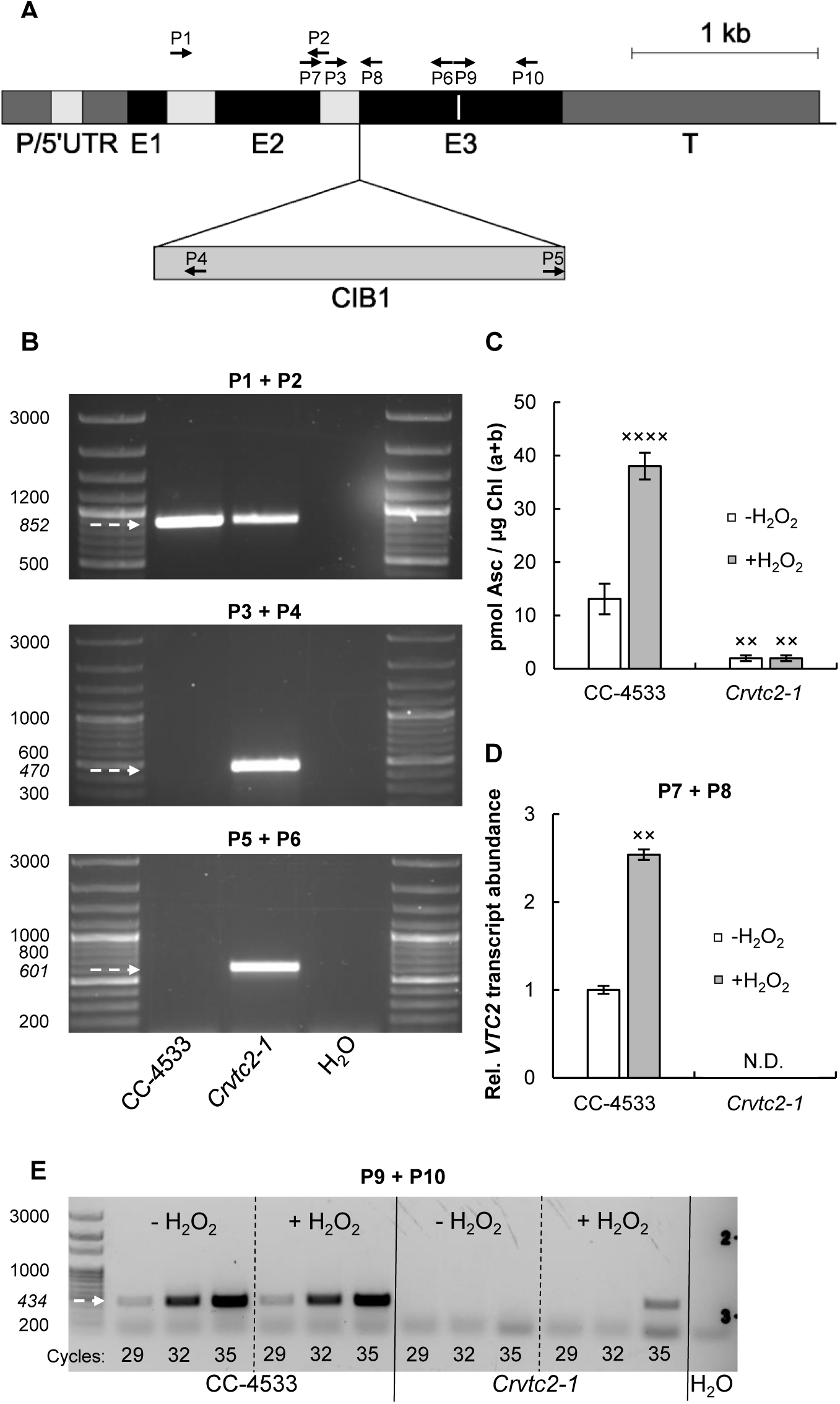
Characterization of an insertional CLiP mutant of *C. reinhardtii* (*LMJ.RY0402.058624*, named *Crvtc2-1*), affected in the *VTC2* gene that encodes GDP-L-galactose phosphorylase. A, Physical map of the *VTC2* gene (obtained from Phytozome v12.1.6) with the CIB1 cassette insertion site in the *Crvtc2-1* mutant. Exons are shown in black, introns in light grey, and promoter/ 5’ UTR and terminator sequences in dark grey. Insertion site of the CIB1 cassette is indicated by triangle and the binding sites of the primers used for genotyping and gene expression analysis of *Crvtc2-1* are shown as black arrows. The sequence encoding the catalytic site of GDP-L-galactose phosphorylase is marked as a white line within Exon 3; B, PCR performed using primers annealing upstream the predicted cassette insertion site in *VTC2* (top panel, using primers P1+P2), and using primers amplifying the 5’ and 3’ genome-cassette junctions (using primers P3+P4 and P5+P6, respectively, middle and bottom panels). The expected sizes are marked with arrows; C, Ascorbate contents of the wild type (CC-4533) and the *Crvtc2-1* mutant grown mixotrophically in TAP medium at moderate light with and without the addition of 1.5 mM H_2_O_2_; D, Transcript levels of *VTC2*, as determined by real-time qRT-PCR in cultures supplemented or not with H_2_O_2_, using primers P7+P8. E, qRT-PCR analysis using primers P9+P10, spanning the sequence that encodes the catalytic site of GDP-L-galactose phosphorylase. The number of PCR cycles is indicated at the bottom of the figure. Data was analyzed by one-way ANOVA followed by Dunnett post-test: × p<0.05, ×× p<0.01, ×××× p<0.0001 compared to the untreated CC-4533 strain.

The site of CIB1 casette integration in the CLiP mutants had been validated by LEAP-Seq method (Li et al., 2016b), and we verified it in the *Crvtc2-1* mutant by PCR (Fig. 1B). Using primers annealing upstream the predicted insertion site in *VTC2*, a specific 852 bp fragment was observed in genomic DNA samples isolated from wild type *C. reinhardtii* cells (CC-4533) and from the *Crvtc2-1* mutant strain (Fig. 1B, top panel); using primers designed to amplify the 5’ and 3’ junction sites of the CIB1 cassette, specific 470 and 601 bp fragments could be detected in the *Crvtc2-1* mutant (Fig. 1B, middle and bottom panels). Sequencing analysis of the PCR amplicons confirmed the predicted insertion of the CIB1 cassette in antisense orientation with its 5’ junction in the third exon of the gene and the 3’ junction reaching to the adjacent intron upstream of exon 3 (Supplemental Fig. S1).

Under moderate light (100 µmole photons m^−2^ s^−1^) and photomixotrophic conditions (growth in Tris-acetate-phosphate (TAP) medium), the wild type strain (CC-4533) had approx. 12 pmol Asc/ µg Chl(a+b) (Fig. 1C), corresponding to about 200 µM cellular Asc concentration (see Kovács et al., 2016 for calculations), and the *Crvtc2-1* mutant had a very low Asc content, only approx. 10% of the wild type. When the cultures were treated with 1.5 mM H_2_O_2_, which had been shown to result in a strong increase in Asc content (Urzica et al., 2012; Vidal-Meireles et al., 2017), the Asc content in the wild type increased approximately three-fold, whereas in the *Crvtc2-1* mutant it did not increase (Fig. 1C). This is in contrast to the *VTC2*-amiRNA lines generated earlier, where H_2_O_2_ treatment resulted in noticeable Asc accumulation (Vidal-Meireles et al., 2017).

Via real time qRT-PCR analysis with primers located upstream and downstream of the insertion site of the CIB1 cassette no *VTC2* transcript could be detected in the *Crvtc2-1* mutant samples, grown under normal growth conditions or treated with H_2_O_2_ (Fig. 1D). Similarly, in qRT-PCR analysis using primers spanning the sequence encoding the catalytic site of *VTC2* (which is located downstream of the CIB1 casette insertion site), no transcript could be detected in the *Crvtc2-1* mutant under normal growth conditions, and only a weak signal could be observed upon 35 PCR cycles in the H_2_O_2_-treated *Crvtc2-1* mutant samples (Fig. 1E).

In order to confirm that the decrease in Asc content is caused by the functional deletion of *VTC2* in the insertional mutant strain, genetic complementation was carried out. To this end, we transformed the *Crvtc2-1* insertional mutant with the coding sequence of *VTC2*, controlled by the constitutive promoter *PsaD*. The plasmid used for transformation included the *APH7”* resistance gene (Fig. 2A), thus the ability to grow on a medium containing hygromycin-B was used as the first screening method for successful transformation.

**Figure 2.**
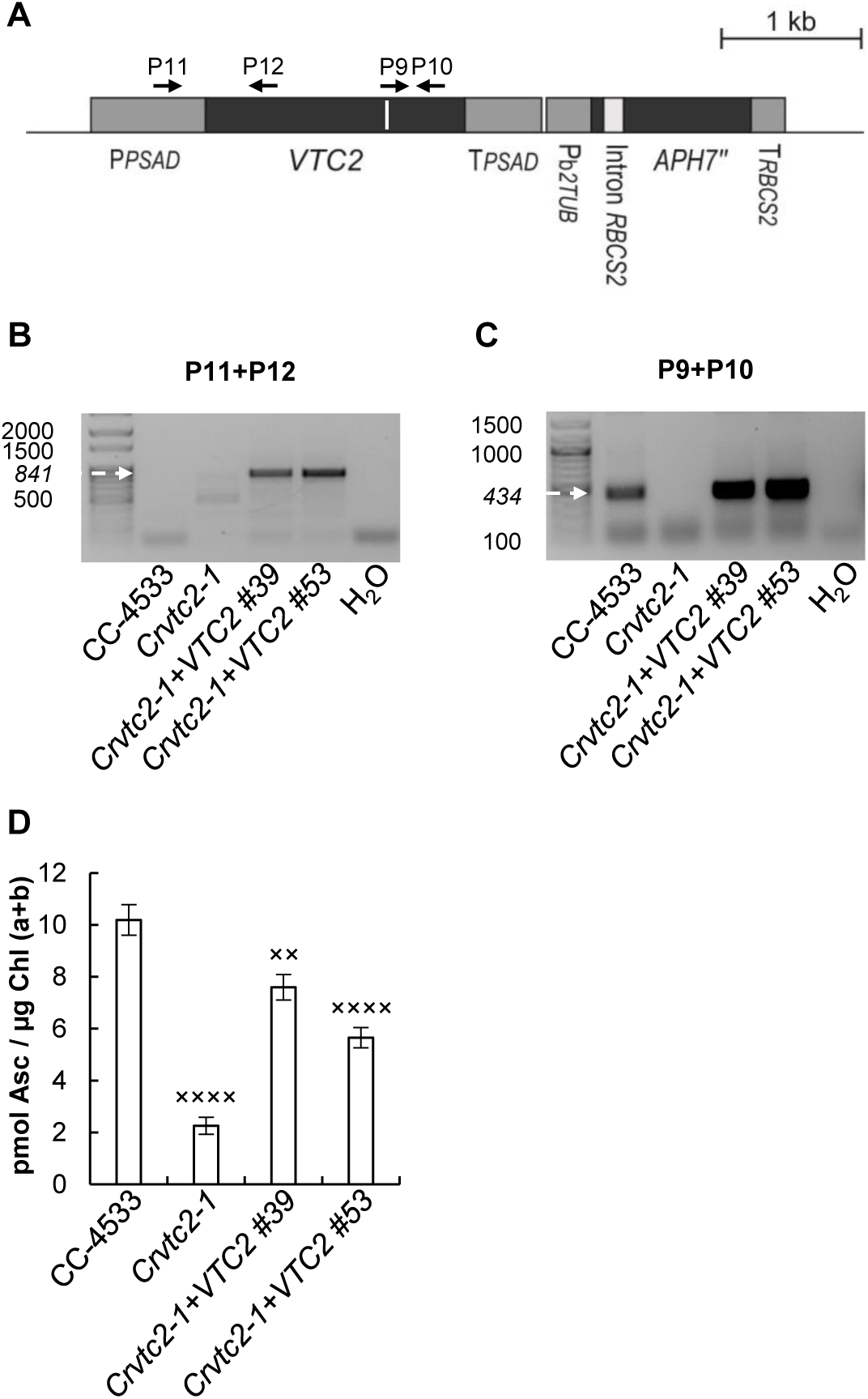
Complementation of the insertional CLiP mutant *LMJ.RY0402.058624*, affected in the *VTC2* gene (named *Crvtc2-1*) with the coding sequence of *VTC2*. A, Physical map of the *Crvtc2-1+VTC2* plasmid containing the coding sequence of *VTC2*, the constitutive promoter *PsaD* and the *APH7”* resistance gene. Exons are shown in black and promoter/ 5’ UTR, terminator sequences in dark grey, and the sequence encoding the catalytic site of GDP-L-galactose phosphorylase is marked as a white line. The binding sites of the primers used below are shown as black arrows; B, PCR performed using primers annealing in the promoter and *VTC2* exon 1 (P11+P12). The expected size is marked with an arrow; C, qRT-PCR performed using primers annealing to the sequence encoding the catalytic site of *VTC2* (P9+P10). The expected size is marked with an arrow; D, Ascorbate contents of CC-4533, the *Crvtc2-1* mutant and the complementation lines *Crvtc2-1+VTC2* grown for 3 days in TAP at 100 µmole photons m^−2^ s^−1^. Data was analyzed by one-way ANOVA followed by Dunnett post-test: × p<0.05, ××× p<0.001, ×××× p<0.0001 compared to the CC-4533 strain. µE stands for µmole photons m^−2^ s^−1^.

The integration of the plasmid in the genome was verified by PCR. Using a forward primer annealing to the *PSAD* promoter region and a reverse primer annealing to the 5’ end of the *VTC2* coding sequence, a specific 841 bp fragment could be amplified in genomic DNA samples isolated from two independent complementation lines of *Crvtc2-1+VTC2* (Fig. 2B). The *VTC2* transcript could be detected in the complemented *Crvtc2-1+VTC2* lines via qRT-PCR analysis with primers spanning the sequence encoding the catalytic site (Fig. 2C). The Asc content of the complementation lines were restored to a great extent (Fig. 2D). The cell volume, the cellular Chl content and Chl*a/b* ratios were moderately increased in the *Crvtc2-1* mutant relatively to the wild type and these parameters were partially restored upon complementation (Supplemental Fig. S2A, B, C).

Regarding the growth phenotypes, no significant difference was observed between the strains when grown in TAP medium at 100 µmole photons m^−2^ s^−1^, whereas the growth of the *Crvtc2-1* mutant was severely inhibited in TAP medium at 530 µmole photons m^−2^ s^−1^, which was restored upon genetic complementation. In HSM medium at 530 µmole photons m^−2^ s^−1^ growth was slow in all genotypes and no significant differences were observed among them (Supplemental Fig. S2D). The Asc content increased two-to three-fold in each strains upon high light treatment, and the Asc content of the *Crvtc2-1* mutant remained at a level of 10-20% relative to the wild type and the complementation lines (Supplemental Fig. S2E).

### The effects of Asc deficiency on NPQ in cultures grown at normal light and photomixotrophic conditions

NPQ includes short-term responses to changes in light intensity, as well as responses that occur over longer periods allowing for acclimation to high light exposure. In *C. reinhardtii*, the levels of the different NPQ components are variable and highly dependent on the growth conditions (Niyogi et al., 1997; Finazzi et al., 2006; Iwai et al., 2007; Peers et al 2009).

In a first set of experiments to assess the effects of Asc deficiency on NPQ, Chlamydomonas strains were cultured in TAP medium, at 100 µmol photons m^−2^ s^−1^. Before the NPQ measurements, cultures were dark-adapted for about 30 min with shaking in order to avoid anaerobiosis; this dark adaptation protocol ensures the relaxation of most NPQ processes and the separation of the NPQ components induced under high light illumination (Roach and Na, 2017). When subjecting the cells to continuous red light of 530 µmol photons m^−2^ s^−1^, a small, rapidly induced NPQ component was induced in the wild type and the Asc-deficient *Crvtc2-1* strain in the first 2 min (Fig. 3A), that we attribute to energy-dependent component (qE). qE is activated by low lumen pH, which occurs for instance during the induction of photosynthesis and upon CO_2_ limitation of the Calvin-Benson-Bassham cycle (Kanazawa and Kramer, 2002; Takizawa et al., 2008). In *C. reinhardtii*, qE formation also requires zeaxanthin or lutein (Ericksson et al., 2015) and it is enhanced by a stress-related LHC protein, LHCSR3, which is strongly expressed when algae are grown at high light (Xue et al., 2015; Peers et al., 2009; Bonente et al., 2011; Chaux et al., 2017). At moderate light (100 µmol photons m^−2^ s^−1^) and photomixotrophic growth conditions, LHCSR3 level was relatively low, particularly in the *Crvtc2-1* strain (Supplemental Fig. S3). The presence of acetate also enables high Calvin-Benson-Bassham cycle activity and a relatively low qE (Johnson and Alric, 2012), in agreement with our findings.

**Figure 3.**
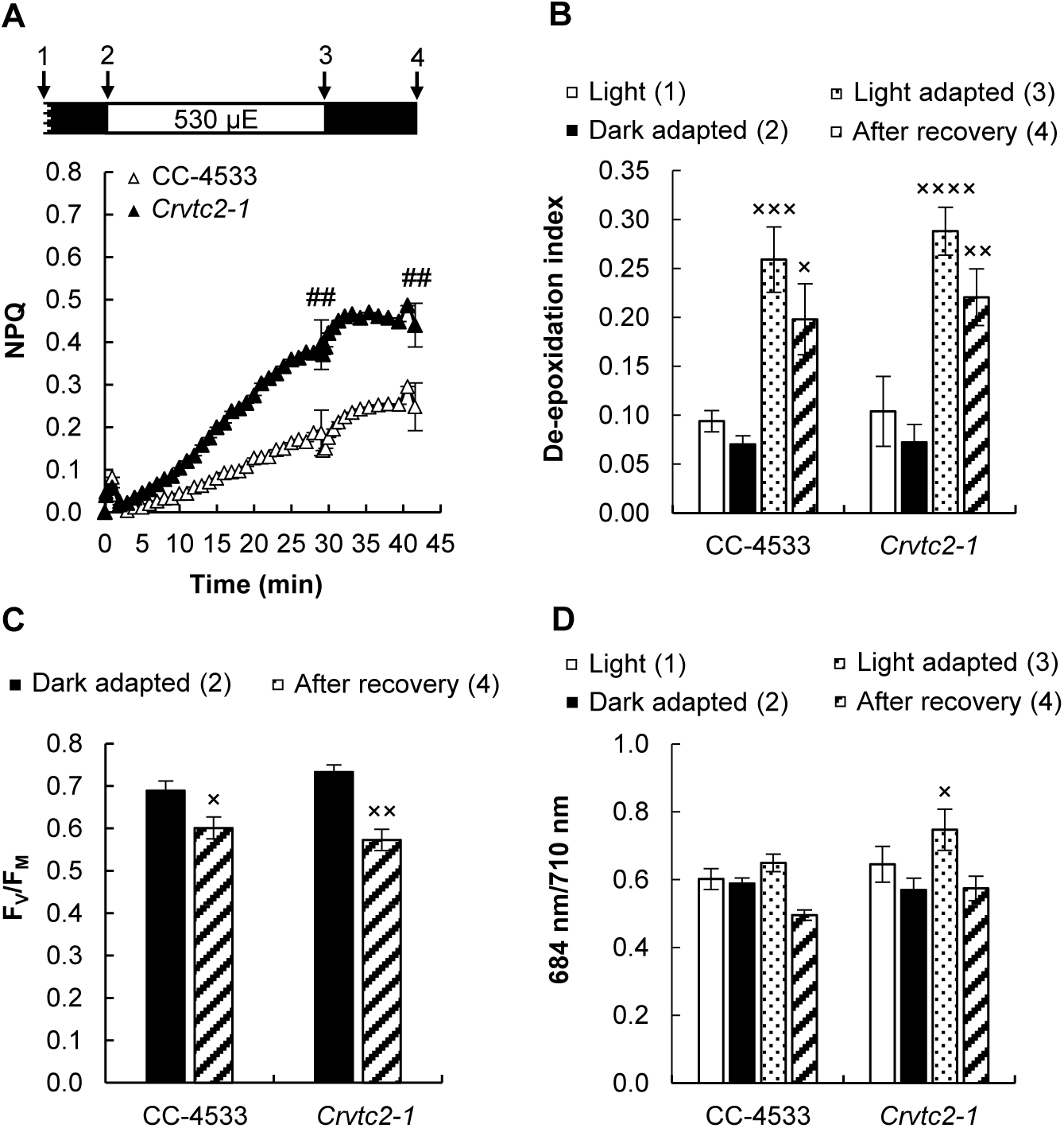
Acclimation to 530 µmole photons m^−2^ s^−1^ of red light followed by recovery in CC-4533 (wild type) and *Crvtc2-1* cultures, grown photomixotrophically in TAP medium at 100 µmole photons m^−2^ s^−1^. A, NPQ kinetics; B, De-epoxidation index; C, F_V_/F_M_ parameter measured after dark adaptation and after recovery from the 530 µmole photons m^−2^ s^−1^ red light; D, 684 nm/ 710 nm ratio of the 77K fluorescence spectra. Samples were collected at the growth light of 100 µmole photons m^−2^ s^−1^, after 30 min of dark-adaptation, at the end of the 30 min light period to 530 µmole photons m^−2^ s^−1^ and 15 min after the cessation of actinic illumination, as indicated by arrows in the scheme in panel A. Data was analyzed by one-way ANOVA followed by Dunnett post-test: ## p<0.01 compared to the CC-4533 strain at the respective time-point; × p<0.05, ×× p<0.01, ××× p<0.001 compared to the dark-adapted CC-4533 strain. µE stands for µmole photons m^−2^ s^−1^.

A slower NPQ component, induced on the timescale of several minutes was also present, which was enhanced in the Asc-deficient *Crvtc2-1* mutant (Fig. 3A) and restored in its complementation strains (Supplemental Fig. S4A). This slow component was enhanced in our previously published *VTC2*-amiRNA line relative to its control strain as well (Supplemental Fig. S4C).

Three components may be responsible for this slowly induced NPQ component (see Ericksson et al., 2015; Allorent et al., 2013): i) zeaxanthin dependent quenching, which may act on NPQ directly (e.g. Holt et al., 2005; Holub et al., 2007; Avenson et al., 2008) or indirectly by controlling the sensitivity of qE to the pH gradient or promoting conformational changes within LHCs (e.g. Johnson et al., 2008; Ruban et al., 2012); ii) State transition-dependent quenching (qT), which may contribute to balancing excitation energy between PSII and PSI via LHCII phosphorylation and antenna dissociation from PSII (Depège, 2003; Lemeille et al., 2009; Ünlü et al., 2014); iii) A slowly relaxing, “photoinhibitory” quenching (qI), associated with photosystem II (PSII) damage or slowly reversible downregulation of PSII representing a continuous form of photoprotection (Adams et al. 2013; Tikkanen et al., 2014).

In order to decipher the origin of the slow NPQ component and to study the possible role of Asc in NPQ, carotenoids were analyzed first, using HPLC. Upon illumination, the de-epoxidation index largely increased (from about 0.05 to 0.25) both in the CC-4533 (wild type) strain and the *Crvtc2-1* mutant, and de-epoxidation only moderately recovered after the cessation of actinic illumination in both strains (Fig. 3B). Violaxanthin, antheraxanthin and zeaxanthin concentrations were essentially the same in the *Crvtc2-1* mutant and in the wild type (Supplemental Fig. S5A, B, C). These results suggest that qZ was partially responsible for the slow NPQ component and that Asc deficiency does not limit the de-epoxidation reaction. We also found that the amounts of β-carotene and lutein were not affected by the lack of Asc and their quantities remained constant during the entire protocol (Supplemental Fig. S5D, E). The F_V_/F_M_ values of dark-adapted cultures and those subjected to high light illumination followed by a recovery period were also very similar, with no major differences between the Asc-deficient mutant and the CC-4533 strain (Fig. 3C).

Since the de-epoxidation ratios were the same in the CC-4533 strain and in the *Crvtc2-1* mutant (Fig. 3B), it is likely that Asc-deficiency does not limit the reaction. In order to completely exclude this possibility, a 16-h dark acclimation experiment was carried out, ensuring undetectably low level of Asc (Fig. 4A). Still, NPQ was induced slowly upon illumination (Fig. 4B), and the de-epoxidation indices were similar than in cultures subjected to relatively short dark adaptation (compare Fig. 4C and Fig. 3B); we note that during a 30-min illumination Asc does not accumulate (Vidal-Meireles et al., 2017).

**Figure 4.**
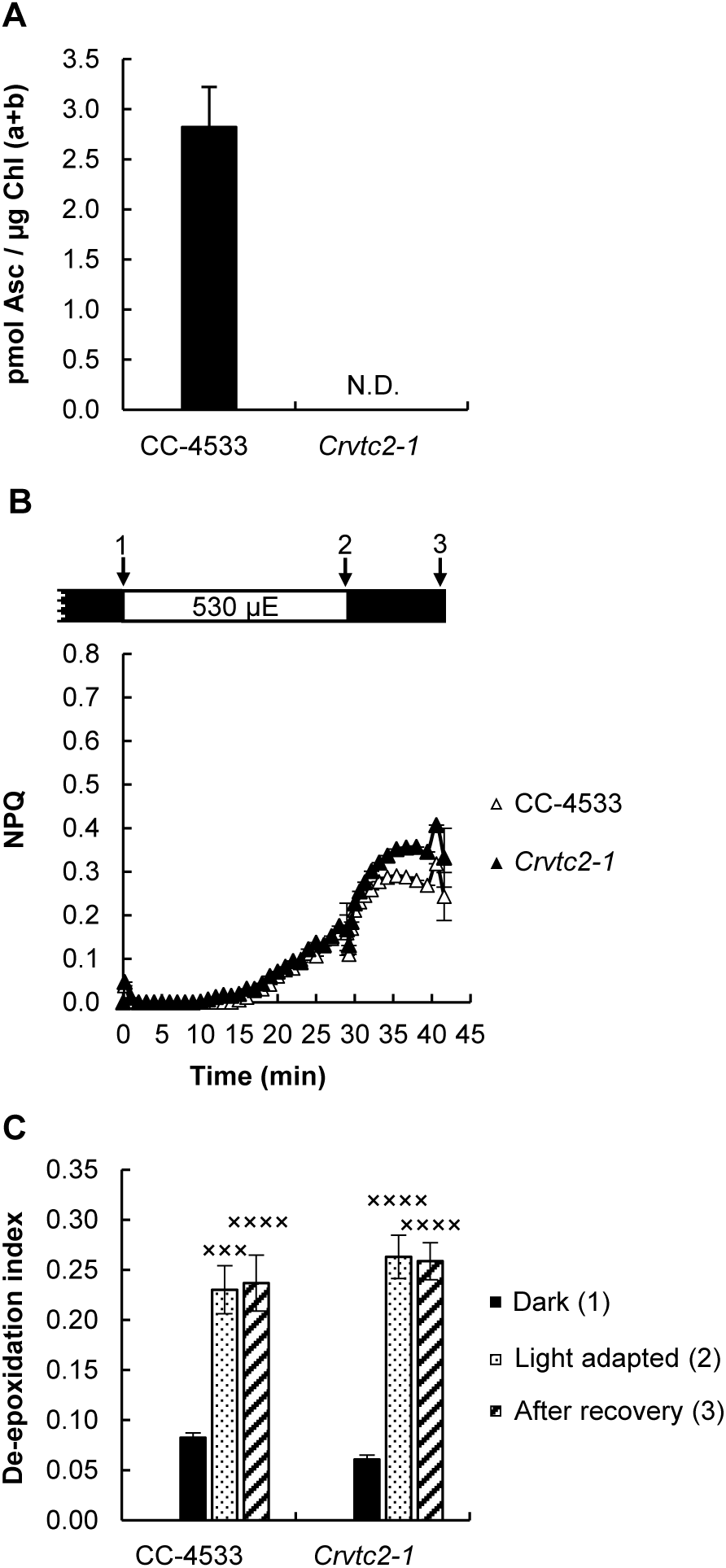
Effects of overnight (16 h) dark acclimation on the CC-4533 and the *Crvtc2-1* (grown in TAP medium at 100 µmole photons m^−2^ s^−1^). A, Ascorbate content after 16 h of dark acclimation; B, NPQ, induced by 530 µmole photons m^−2^ s^−1^ of red light after overnight dark acclimation; C, de-epoxidation index, determined in the overnight dark-acclimated cultures, after strong red-light illumination and following recovery. Data was analyzed by one-way ANOVA followed by Dunnett post-test: ××× p<0.001, ×××× p<0.0001 compared to the dark-acclimated CC-4533 strain. µE stands for µmole photons m^−2^ s^−1^.

The large increase in the de-epoxidation index upon illumination suggests that qZ is at least partially responsible for the slow NPQ component. However, we also observed that the slow NPQ component was larger in the *Crvtc2-1* mutant than in the wild type, whereas the de-epoxidation ratios were the same (Fig. 3A, B). In addition to qZ, qT and qI mechanisms may also contribute to the slow component and they may differ between the wild type and the *Crvtc2-1* mutant. The possible contribution of qT was studied by measuring 77K fluorescence spectra: Upon illumination with 530 µmole photons m^−2^s^−1^ red light, the 684 nm/710 nm ratio remained unaltered in the wild type and increased slightly in the Asc-deficient mutant (Fig. 3D). Transition from state I to state II would decrease the 684/710 nm ratio, therefore, in our cultures grown at moderate light in TAP medium and subjected to strong red illumination during the fluorescence measurement, qT is unlikely to contribute to NPQ induction. On the other hand, when the actinic illumination was switched off, the 684/710 nm ratio decreased moderately, reflecting the occurrence of state I to state II transition in the dark.

To further study the effect of state transition in the induction and relaxation of NPQ, a state transition mutant, called *stt7-9* (Depège et al., 2003) was employed. NPQ was induced during illumination in the *stt7-9* mutant to a similar extent as in the *Crvtc2-1* mutant (albeit with rather different kinetics), which coincided with a strong zeaxanthin accumulation (Supplemental Fig. S6B); this indicates that transition to state II did not play a role in the formation of NPQ under the present experimental conditions. On the other hand, upon the cessation of actinic illumination, there was a rapid NPQ relaxation in the *stt7-9* mutant, showing that transition to state II occurs in the wild types and the *Crvtc2-1* mutant in the dark, probably masking the relaxation of the other NPQ components.

As a next step, the effect of oxidative stress, known to enhance NPQ (Roach and Na, 2017), was tested by employing H_2_O_2_ and catalase treatments on the *Crvtc2-1* mutant and its wild type. Fig. 5A and B show that upon the addition of 1.5 mM H_2_O_2_, the slow NPQ component increased remarkably in both strains, without altering the de-epoxidation level (Fig. 5C). When 5 µg/ml catalase was added, NPQ was only slightly affected in the wild type (Fig. 5D), whereas it significantly decreased in the Asc-deficient mutant (Fig. 5E). These data suggest that H_2_O_2_ accumulated upon strong illumination in the Asc-deficient mutant, resulting in enhanced NPQ. On the other hand, the F_V_/F_M_ value, an indicator of photosynthetic efficiency, did recover following illumination and to a similar extent in the wild type and the *Crvtc2-1* strain (Fig. 3C), thus photosynthetic reaction centers did not get severely inhibited.

**Figure 5.**
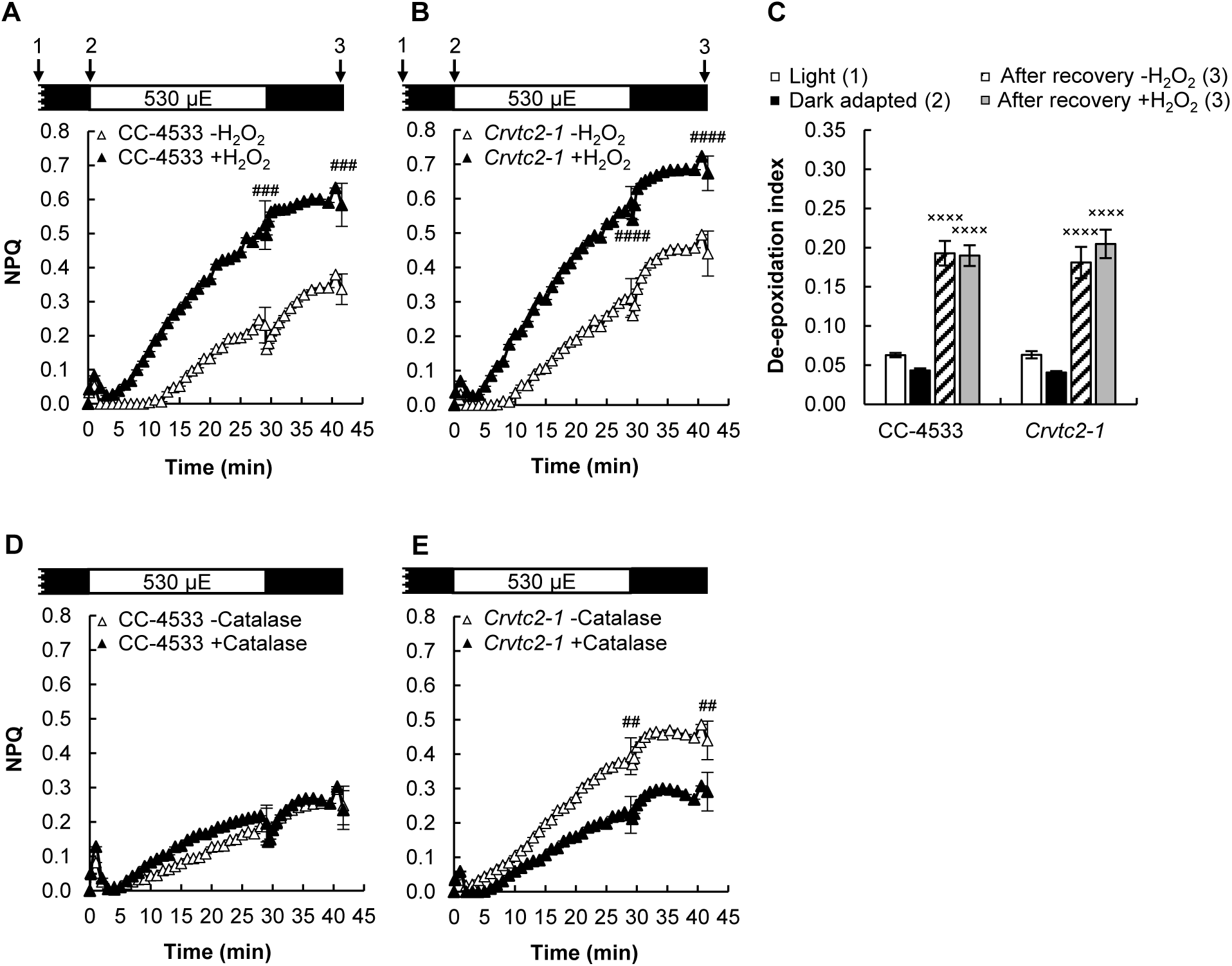
The effects of H_2_O_2_ and catalase on NPQ, induced by strong red light (530 µmole photons m^−2^ s^−1^) in the wild type (CC-4533) and the *Crvtc2-1* mutant grown in photomixotrophic conditions in TAP medium at 100 µmole photons m^−2^ s^−1^. A, The effect of 1.5 mM H_2_O_2_ on NPQ induction in the CC-4533 strain; B, the effect of 1.5 mM H_2_O_2_ on NPQ induction in the *Crvtc2-1* mutant; C, the effect of H_2_O_2_ addition on de-epoxidation; D, the effect of catalase on NPQ induction in the CC-4533 strain; E, the effect of catalase on NPQ induction in the *Crvtc2-1* mutant. Samples were collected at the time points indicated by arrows in the schemes in panels A and B. Data was analyzed by one-way ANOVA followed by Dunnett post-test: ## p<0.01, ### p<0.001, #### p<0.0001 compared to the untreated CC-4533 culture at the respective time-point; × p<0.05, ×× p<0.01, ××× p<0.001 compared to the dark-adapted CC-4533 strain. µE stands for µmole photons m^−2^ s^−1^.

For comparison, the *npq1* mutant, lacking the CVDE enzyme, thus unable to perform violaxanthin de-epoxidation (Niyogi et al., 1997), was also tested. Upon illumination with 530 µmol photons m^−2^ s^−1^, this strain developed a large NPQ (Fig. 6A), which was accompanied by irreversible decrease of F_V_/F_M_ and loss of Chl and β-carotene relative to its wild type (137a) strain (Fig. 6B, C, D). 77 K fluorescence recordings showed no changes in the 684nm/710nm ratio (Fig. 6E), thus the large NPQ component could be unambiguously attributed to photoinhibitory qI. Interestingly, the Asc concentration in the *npq1* mutant is very high compared to the other strains (Fig. 6F), probably to compensate for the lack of CVDE and zeaxanthin in ROS management (Baroli et al., 2003). Thus, the experiments on the *npq1* mutant corroborate the importance of CVDE in strong illumination.

**Figure 6.**
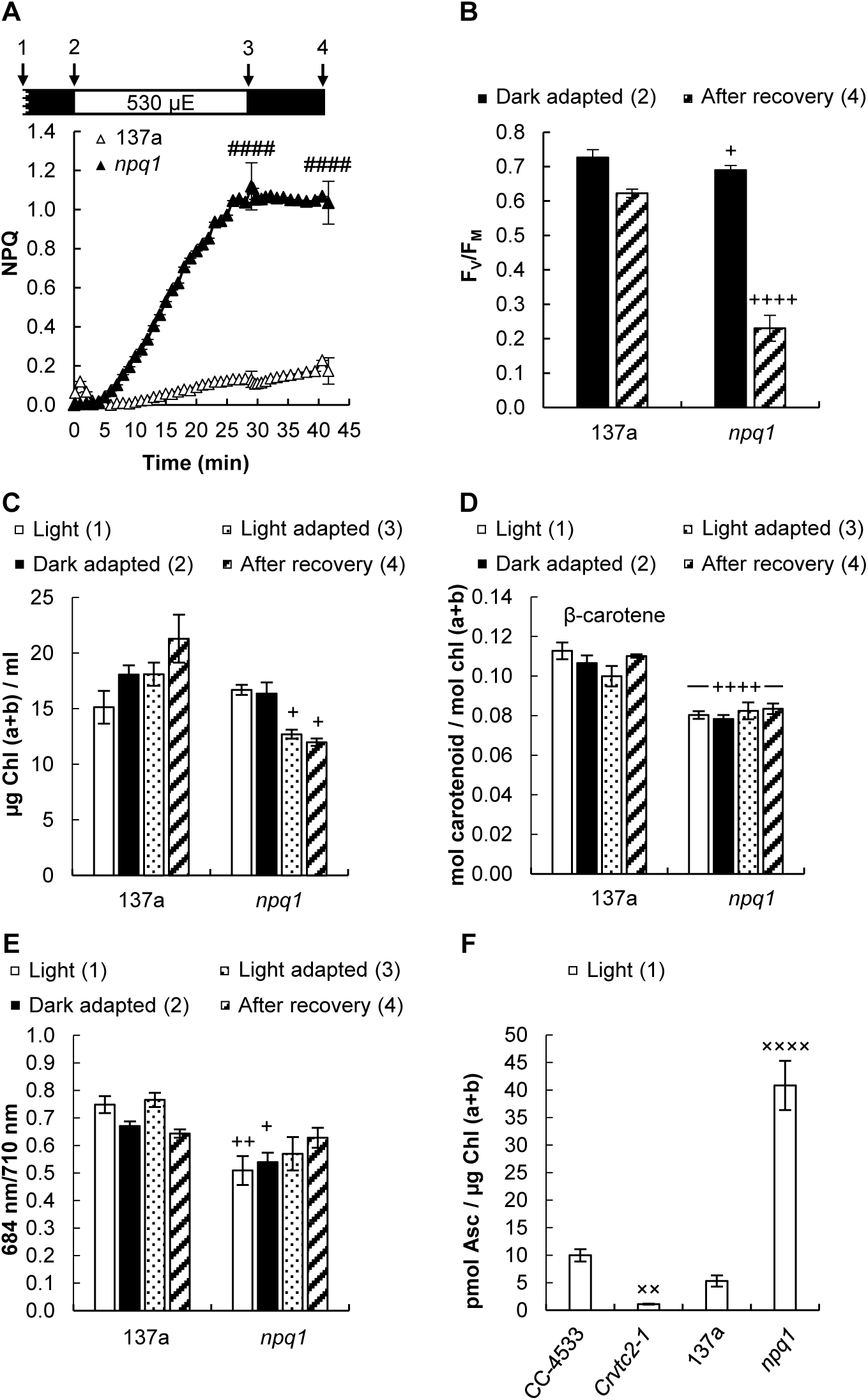
Effects of strong red light (530 µmole photons m^−2^ s^−1^) on the 137a (wild type) and the *npq1* mutant of *C. reinhardtii* grown in TAP medium at 100 µmole photons m^−2^ s^−1^. A, NPQ induced by 530 µmole photons m^−2^ s^−1^ of red light followed by a recovery phase; B, F_V_/F_M_ value, determined before the strong red light illumination and after the recovery phase; C, Chl(a+b) content of the cultures determined before, during and after the strong red light illumination; D, β-carotene content measured before, during and after the strong red light illumination; E, 684 nm/ 710 nm ratio of the 77K fluorescence spectra determined before, during and after the strong red light illumination; F, Ascorbate contents of the *npq1* and *Crvtc2-1* mutants and the CC-4533 and 137a wild type strains. Samples were collected at the time points indicated by arrows in the scheme in panel A. Data was analyzed by one-way ANOVA followed by Dunnett post-test: #### p<0.0001 compared to the 137a strain at the respective time-point; + p<0.05, ++ p<0.01, ++++ p<0.0001 compared to the dark-adapted 137a strain; ×× p<0.01, ×××× p<0.0001 compared to the CC-4533 strain. µE stands for µmole photons m^−2^ s^−1^.

### The effects of Asc deficiency on NPQ in cultures grown under photoautotrophic conditions at high and moderate light

When the cultures were grown under photoautotrophic conditions without CO_2_ supplementation under strong white light (530 µmole photons m^−2^s^−1^), which was similar in intensity used for NPQ induction measurements, qE reached relatively high values (about 1.0) both in the wild type and the Asc-deficient CLiP mutant (Fig. 7A). In the *VTC2*-amiRNA line, the qE component was enhanced relative to its empty vector control (Supplemental Fig. S4D). These results show that Asc is not required for the formation of the qE component. The qE phase was followed by a slower one, which was enhanced both in the *Crvtc2-1* mutant and the *VTC2* amiRNA line relative to their control strains.

**Figure 7.**
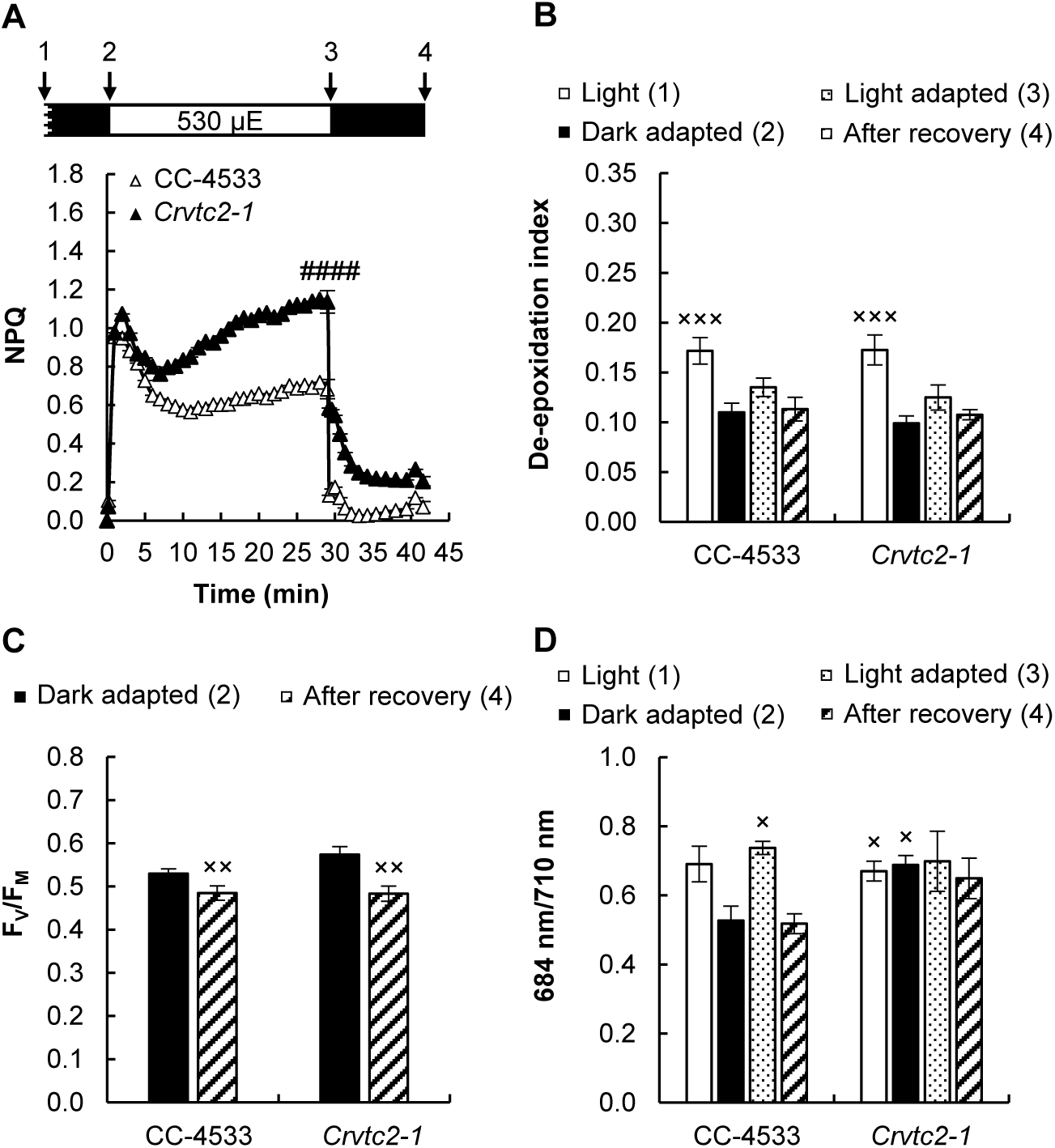
Acclimation to 530 µmole photons m^−2^ s^−1^ of red light followed by recovery in CC-4533 and *Crvtc2-1* cultures, grown photoautotrophically in HSM medium at 530 µmole photons m^−2^ s^−1^. A, NPQ kinetics; B, De-epoxidation index; C, F_V_/F_M_ parameter measured after dark adaptation and after recovery from the 530 µmole photons m^−2^ s^−1^ red light illumination; D, 684 nm/ 710 nm ratio of the 77K fluorescence spectra. The samples were collected at the growth light of 530 µmole photons m^−2^ s^−1^, after 30 min of dark adaptation, at the end of the 30 min red light illumination, and 12 min after the cessation of actinic illumination, as indicated in the scheme in panel A. Data was analyzed by one-way ANOVA followed by Dunnett post-test: #### p<0.0001 compared to the CC-4533 strain at the respective time-point; × p<0.05, ×× p<0.01, ××× p<0.001 compared to the dark-adapted CC-4533 strain. µE stands for µmole photons m^−2^ s^−1^.

During illumination, the de-epoxidation index changed only marginally, and it was essentially the same in the wild type and in the Asc-deficient strain (about 0.1, Fig. 7B). The F_V_/F_M_ value was also unaffected in the *Crvtc2-1* mutant relative to its wild type before or after the illumination with strong red light (Fig. 7C). The 684nm/710nm ratio of the 77 K spectra remained constant in the Asc-deficient mutant (Fig. 7D). The violaxanthin, antheraxanthin, zeaxanthin and lutein contents did not decrease upon illumination with intense red light in either strain (Supplemental Fig. S7), only the total amount of β-carotene was slightly lower in the Asc-deficient mutant (Supplemental Fig. S7D). We also observed that under high light growth conditions, the amount of photosynthetic complexes (namely PsbA, CP43, PSBO, PsaA, LHCSR3, PetB and RbcL) were essentially the same in the *Crvtc2-1* mutant and in the wild type, as detected by western blot analysis on equal chl basis (Supplemental Fig. S3).

Treatments with 1.5 mM H_2_O_2_ led to alteration of the NPQ kinetics and a slower relaxation in both strains (Fig. 8A, B). In the *Crvtc2-1* mutant, catalase treatment resulted in a strong decrease of qE and the slow NPQ component (Fig. 8C, D). These results show that under photoautotrophic and high light conditions, Asc-deficiency does not limit qE or qZ but may lead to the occurrence of oxidative stress and thereby to increased qI.

**Figure 8.**
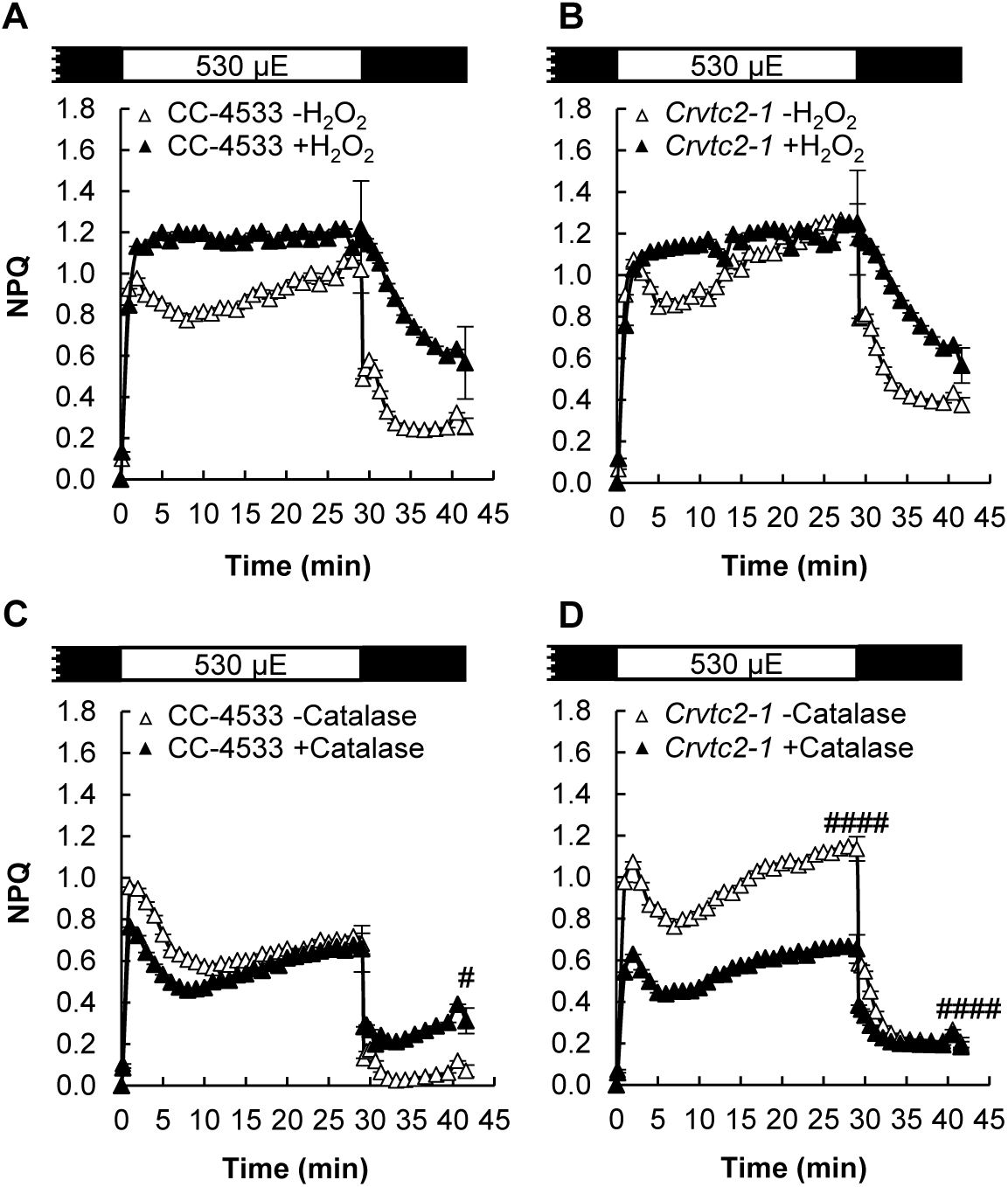
The effects of H_2_O_2_ and catalase on NPQ induced by strong red light (530 µmole photons m^−2^ s^−1^) in the wild type (CC-4533) and *Crvtc2-1* mutant strains grown photoautotrophically in HSM medium at 530 µmole photons m^−2^ s^−1^. A, The effect of 1.5 mM H_2_O_2_ on NPQ induction in the CC-4533 strain; B, the effect of 1.5 mM H_2_O_2_ on NPQ induction in the *Crvtc2-1* mutant; C, the effect of catalase on NPQ induction in the CC-4533 strain; D, the effect of catalase on NPQ induction in the *Crvtc2-1* mutant. Data was analyzed by one-way ANOVA followed by Dunnett post-test: # p<0.05, #### p<0.0001 compared to the untreated CC-4533 culture at the respective time-point. µE stands for µmole photons m^−2^ s^−1^.

Subjecting the cells in HSM medium in moderate light (100 µmole photons m^−2^s^−1^) resulted in similar effects in terms of qE, de-epoxidation, the 684/710 nm ratio of the 77 K spectra and H_2_O_2_ and catalase responsiveness (Supplemental Fig. S8).

## Discussion

### The Crvtc2-1 CLiP mutant possesses a low Asc content without major changes in the phenotype

*VTC2* encodes GDP-L-galactose phosphorylase, an essential and highly regulated enzyme of Asc biosynthesis both in vascular plants and in green algae (Urzica et al., 2012; Vidal-Meireles et al., 2017) and downregulating *VTC2* via the amiRNA technique results in Asc deficiency (Vidal-Meireles et al., 2017). For the present study, we identified and genetically complemented a *VTC2* mutant in the CLiP collection that carries a single insertion in the *VTC2* gene (Fig. 1 and Fig. 2). The *Crvtc2-1* mutant possesses about 10% Asc relative to its wild type strain CC-4533 under normal growth conditions, its Asc content is unaltered upon H_2_O_2_ treatment and remains below 20% of the wild type under high light conditions; in addition, by employing overnight dark acclimation, the Asc concentration of the *Crvtc2-1* mutant strongly decreased (Fig. 4).

The *Crvtc2-1* mutant is very likely to be a knockout for *VTC2* as no transcript accumulation could be detected performing qRT-PCR with primers annealing downstream of the CIB cassette insertion site and spanning the sequence encoding the catalytic site of GDP-L-galactose phosphorylase. Only when the cultures were treated with H_2_O_2_ and when a high PCR cycle number was used (Fig. 1E) a faint band could be observed in the gel. It is very unlikely that a functional truncated GDP-L-galactose phosphorylase is present in the mutant, but the observation that the *Crvtc2-1* strain still contains 10-20% Asc relative to its parent strain suggests that some phosphorolysis of GDP-L-galactose could be carried out by another enzyme ensuring minor amount of Asc. We note that in Arabidopsis *VTC2* has a lowly expressed homologue, *VTC5*, and knocking out both of them results in seedling lethality (Dowdle et al., 2007). In Chlamydomonas, no homologue of *VTC2* has been identified (Urzica et al., 2012). On the other hand, it is possible that GDP-L-galactose is degraded hydrolytically (with L-galactose-1-P and GMP as products) leading to minor Asc production in the *VTC2* mutant. An alternative Asc biosynthesis pathway may also exist in Chlamydomonas, although homologues of enzymes possibly involved in alternative Asc biosynthesis pathways in vascular plants could not be found in Chlamydomonas (Urzica et al., 2012; Wheeler et al., 2015).

In spite of the very low Asc content of the *Crvtc2-1* mutant, the phenotype was only moderately altered. The *Crvtc2-1* mutant has the same growth rate than the wild type and the complementation lines at moderate light conditions in TAP medium (Supplemental Fig. S2). The amounts of various photosynthetic subunits were similar in the Asc-deficient *Crvtc2-1* mutant than in the wild type strain CC-4533 in TAP medium at moderate light and also in HSM medium both at moderate and high light (Supplemental Figure S3). Unexpectedly, the amount of the photoprotective LHCSR3 protein was reduced in the *Crvtc2-1* mutant in TAP medium at moderate light and it was at the same level than in the wild type when grown in HSM medium both at medium and high light. The amounts of carotenoids are unchanged in the *Crvtc2-1* line in cultures grown at normal light, whereas at high light in HSM medium, the amount of β-carotene is slightly reduced (Supplemental Fig. S5 and S7). The *Crvtc2-1* line had a slightly higher Chl content than its wild type and cell size was also moderately increased (Fig. 2). A marked characteristics of the *Crvtc2-1* mutants was that it was unable to grow at high light in TAP medium (Supplemental Fig. S2).

In our previously published *VTC2*-amiRNA line Asc deficiency led to more severe alterations in the phenotype than it was observed in the *Crvtc2-1* mutant (Vidal-Meireles et al., 2017). The reason behind this remains to be elucidated, although the cell wall deficiency of the cw15-325 line (the parent strain of the *VTC2*-amiRNA line) and thereby its increased stress sensitivity (Voigt and Münzner, 1994) may explain the differences between the *VTC2*-amiRNA and the *Crvtc2-1* insertional mutant strains.

### The effects of Asc deficiency on the qE component of NPQ

*C. reinhardtii* uses various photoacclimation strategies which strongly depend on the carbon availability and the trophic status of the cells (Polukhina et al., 2016). The fast rise in NPQ (qE) is enhanced upon growth at high light and low CO_2_ that is enabled by a high expression of LHCSR3 (e.g. Peers et al., 2009). It has been observed that under photomixotrophic conditions at normal light, the expression of the LHCSR proteins is very low and qE is minor; in addition, the deepoxidation state is also known to vary with the growth light (Polukhina et al., 2016). Therefore, in order to study the role of Asc in the different NPQ parameters, we subjected the cultures both to moderate and high light, photomixotrophic and photoautotrophic conditions. As shown by our results and the discussion below, by these means we managed to distinguish between qE, qZ, and qI, and only qT could not be studied in detail.

Rapid response to changes in light intensity and dissipation of excess light energy are particularly important when the activity of the Calvin-Benson-Bassham cycle is limiting, in order to avoid a potentially deleterious buildup of excessive ΔpH (Kanazawa and Kramer, 2002; Takizawa et al., 2008). In agreement with the literature (Xue et al., 2015), at normal light and photomixotrophic conditions, the rapidly inducible qE was a minor component and the relative amount of the LHCSR3 protein, essential for qE development was low (Supplemental Fig. S3). When Chlamydomonas cultures were grown at photoautotrophic, possibly CO_2_-limiting conditions, the amplitude of qE largely increased both at moderate and high light (Supplemental Fig S8 and Fig. 8, respectively) enabled by the accumulation of LHCSR3 (Supplemental Fig. S3) and possibly by other factors.

Our results on the *Crvtc2-1* line show that qE is not limited by Asc deficiency neither at low light nor at high light conditions, nor under photomixotrophic and photoautotrophic conditions; in the *VTC2*-amiRNA line, qE was even enhanced relative to its control line (Supplemental Fig. S4B).

### The effects of Asc deficiency on the slow NPQ components

In Chlamydomonas, a slowly induced NPQ component, with several underlying mechanisms, may also be induced. When CC-4533 cultures were grown at normal light in TAP medium and subjected to strong red light, the major slow component was probably qZ, as shown by the large increase in de-epoxidation (Fig. 3) and by the loss of NPQ induction in the *npq1* mutant (Fig. 6). In cultures grown at high light and photoautotrophic conditions, de-epoxidation was minor upon light adaptation with strong red light (Fig. 7) and it was intermediate when the cultures were grown photoautotrophically at moderate light (Supplemental Fig. S8). De-epoxidation was equal in the *Crvtc2-1* mutant and the wild type in all growth conditions and also upon overnight dark acclimation that led to undetectably low Asc content in the *Crvtc2-1* mutant. These results clearly show that Asc deficiency is not limiting qZ, thus Asc is not used as a reductant by CrCVDE.

The xanthophyll cycle, in which violaxanthin is converted into zeaxanthin during light acclimation, is ubiquitous among green algae, mosses and plants, with exception of Bryopsidales, a monophyletic branch of the Ulvophyceaes in which NPQ is neither related to a pH-dependent mechanism, nor modulated by the activity of the xanthophyll cycle (Christa et al., 2017). It has also been shown that among green alga species, large variations exist in the activity of xanthophyll cycle and in its overall contribution to NPQ, which seems to depend on the environmental selection pressure and less on the phylogeny (Quaas et al., 2015). In mosses, the xanthophyll cycle was shown to significantly contribute to excess energy dissipation upon stress conditions (e.g. Azzabi et al., 2012).

The de-epoxidation reaction itself is catalyzed by distinct enzymes in vascular plants and in Chlorophyceae, including Chlamydomonas (Li et al., 2016a). Plant-type VDE is associated with the thylakoid membrane on the luminal side, where it catalyzes the de-epoxidation reaction of violaxanthin, found in free lipid phase and it uses Asc as a reductant (Hager and Holocher, 1994; Arnoux et al., 2009). CVDE is located on the stromal side of the thylakoid membrane, and, just like higher plant VDE, it also requires a build-up of ΔpH for its activity (Li et al., 2016a). CVDE is related to lycopene cyclases of photosynthetic bacteria, called CruA and CruP (Li et al., 2016a, Bradbury et al., 2012). We have demonstrated in this paper that Asc is not required for the de-epoxidation reaction, and, in general, for qZ in Chlamydomonas.

Green algae contain very small amounts of Asc relative to vascular plants, and, as stated above, effective de-epoxidation is achieved by an enzyme that does not require Asc as a reductant. Interestingly, brown algae, which produce minor amounts of Asc as well, diadinoxanthin de-epoxidase uses Asc as a reductant with much higher affinity for Asc than plant-type VDE, in combination with a shift of its pH optimum towards lower values enabling efficient de-epoxidation (Grouneva et al., 2006). Mosses have plant-type VDE enzymes (Pinnola et al., 2013), which probably require Asc as a reductant. Since mosses contain approx. ten times less Asc compared to vascular plants (Gest et al., 2013), it remains to be explored how this low amount of Asc allows a rapid and intensive development of NPQ, characteric of mosses (e.g. Marschall and Proctor, 2004).

In Chlamydomonas, light and O_2_ availability-dependent state transition (qT), involving major reorganizations of LHCs, also modulate NPQ (Depège, 2003; Lemeille et al., 2009; Ünlü et al., 2014). Under our experimental conditions, ensuring aeration during both the dark and the light adaptation, and using strong red light as actinic light, no decrease occurred in the 685/710 nm ratio of the 77 K fluorescence spectra, suggesting that state I to state II transition did not affect the NPQ induction in the wild type nor in the Asc-deficient strains. The *stt7-9* mutant, unable to perform state transition, did not show decreased NPQ (Supplemental Fig. S6), which would be expected if state transition would constitute a major form of NPQ under our experimental conditions. However, our data do not exclude the possibility that Asc may participate in state transition under conditions favoring its occurrence.

A fourth, and rather complex component of NPQ is qI, possibly with several underlying mechanisms involved (Adams et al., 2013; Tikkanen et al., 2014). We observed both under photomixotrophic conditions at moderate light and under photoautotrophic conditions that the slow NPQ component was enhanced in the *Crvtc2-1* mutant upon illumination with strong red light, which, however, was not attributable to qZ or to qT. Ascorbate deficiency is accompanied by an increase in the intracellular H_2_O_2_ content in Chlamydomonas (Vidal-Meireles et al., 2017) and ROS are known to enhance NPQ via several mechanisms (Roach and Na, 2017). Using H_2_O_2_ and catalase treatments (Fig. 5 and 8, Supplemental Fig. S8), we clearly show that ROS formation is involved in the slowly induced NPQ component in the Asc-deficient strain that can be interpreted as qI. In the wild type, the contribution of qI to NPQ was probably minor under our experimental conditions, since catalase treatment did not diminish NPQ formation (Fig. 5D).

In conclusion, our results reveal fundamental differences between vascular plants and Chlamydomonas regarding the role of Asc in NPQ. Whereas in vascular plants, the most prominent role of Asc is to be a reductant of VDE, in Chlamydomonas, it is pertinent in preventing ROS formation that would lead to photoinhibitory quenching mechanisms.

## Materials and Methods

### Algal strains and growth conditions

*Chlamydomonas reinhardtii* strains CC-4533 (designated as wild type) and *LMJ.RY0402.058624* (designated as *Crvtc2-1*) were obtained from the CLiP library (Li et al., 2016b). The 137a (CC-125) strain and the *npq1* (CC-4100) mutant were obtained from the Chlamydomonas Resource Center (https://www.chlamycollection.org/). The *ARG7* complemented strain *cw15-412* (provided by Dr Michael Schroda (Technische Universität Kaiserslautern, Germany)) was used as control for the *stt7-9* mutant (Depège et al., 2003). The *VTC2*-amiRNA strain and its control *EV2* strain are described in Vidal-Meireles et al. (2017).

The synthetic coding sequence of *VTC2* including a 38 bp-long upstream sequence homologous to the *PSAD* 5’UTR with the BsmI restriction enzyme recognition site was ordered from Genecust (www.genecust.com). The *VTC2* insert was ligated as BsmI/EcoRI fragment into the similarily digested pJR39 (Neupert et al., 2009) vector, resulting in vector pJR112. Finally, pJR112 was digested with BsmI and SmaI and the *VTC2*-containing BsmI/SmaI fragment was ligated to the similarily digested pJR91 vector that carries the *APH7*” resistance marker for selection on hygromycin-B. Transformation of the *Crvtc2-1* mutant strain was done via electroporation in a Bio-Rad GenePulser Xcell™ instrument, at 1000 V, with 10 F capacitance and infinite resistance usinga 4-mm gap cuvette. The cells were plated onto selective agar plates (TAP + 10 µg/ml hygromycin-B) and colonies were picked after 10 days of growth under moderate light (80 µmole photons m^−2^ s^−1^).

Chlamydomonas pre-cultures were grown in 50-ml Erlenmeyer flasks in Tris-acetate phosphate (TAP) medium for three days at 22°C and 100 µmole photons m^−2^ s^−1^ on a rotatory shaker. Following this phase, cultures were grown in 100-ml Erlenmeyer flasks photomixotrophically (in TAP medium) or photoautotrophically (in high salt minimal (HSM) medium) at 22°C at 100 or 530 µmole photons m^−2^ s^−1^ for two additional days. The initial cell density was set to 1 million cells/ml.

### DNA Isolation and PCR

Total genomic DNA from *C. reinhardtii* strains CC-4533 and *Crvt2-1* (*LMJ.RY0402.058624*) was extracted according to published protocols (Barahimipour et al., 2015; Schroda et al., 2001), and 1 µl of the extracted DNA were used as template for the PCR assays, using the GoTaq DNA polymerase (Promega GmbH).

To confirm the CIB1 insertion site in the *Crvt2-1* strain, PCR assays were conducted using gene specific primers that anneal upstream and downstream of the predicted insertion site of the cassette as well as primers specific for the 5’ and 3’ end of the CIB cassette. Primers 1 (5′-TGATGGCCAAGGGCTTAGTG-3′) and 2 (5’-CCGCAAACACCATGCAATCT-3′) amplified the region of the gene upstream the predicted site of CIB1 cassette insertion (control amplicon with an expected size of 852 bp), primers 3 (5′-AGATTGCATGGTGTTTGCGG-3′) and 4 (5′-CAGGCCATGTGAGAGTTTGCC-3′) amplified the 3’ junction site of the CIB1 cassette (amplicon with an expected size of 470 bp), and primers 5 (5′-GCACCAATCATGTCAAGCCT-3′) and 6 (5′-TGTTGTAGCCCACGCGGAAG-3′) amplified the 5’ junction site of the cassette (amplicon with an expected size of 601 bp). The primers 11 (5′-GCTCTTGACTCGTTGTGCATTCTAG-3′) and 12 (5’-CACTGAGACACGTCGTACCTG - 3′) amplified the 3’ junction site of the *PsaD* promoter with the *VTC2* gene in the plasmid used for complementation (amplicon with an expected size of 841 bp).

### Analyses of gene expression by qRT-PCR

Sample collection and RNA isolation was performed as in Vidal-Meireles et al., (2017). The primer pairs for the *VTC2* gene and the reference genes (*bTub2* - Cre12.g549550, *actin* - Cre13.g603700, *UBQ* - XP_001694320) used in real time qRT-PCR were published earlier in Vidal-Meireles et al. (2017). The annealing sites of the primers for analyzing *VTC2* expression are indicated as primers 7 and 8 in Fig. 1. Primers 9 (5′- AACCACCTGCACTTCCACGCTTAC-3′) and 10 (5’-TGCCCCGCAATCTCAAACGATG-3′) spanned the sequence encoding the catalytic site of *VTC2* (amplicon with an expected size of 434 bp).

The real time qRT-PCR data are presented as fold-change in mRNA transcript abundance of the *VTC2* gene, normalized to the average of the three reference genes, and relative to the untreated CC-4533 strain. Real-time qRT-PCR analysis was carried out with three technical replicates for each sample and three biological replicates were measured; the standard error was calculated based on the range of fold-change by calculating the minimum and the maximum of the fold-change using the standard deviations of ΔΔCt.

### Determination of cell size, cell density, chlorophyll, Asc and carotenoid contents

The cell density was determined by a Scepter™ 2.0 hand-held cell counter (Millipore), as described in Vidal-Meireles et al., (2017).

The Chl content was determined according to Porra (1989) and the Asc content was determined as in Kovács et al., (2016).

For carotenoid content determination, liquid culture containing 30 µg Chl(a+b)/ml was filtered onto a Whatman glass microfibre filter (GF/C) and frozen in liquid N_2_ at different time points in the NPQ induction protocol. The pigments were extracted by re-suspending the cells in 500 µl of ice-cold acetone. After re-suspension, the samples were incubated in the dark for 30 min. This was followed by centrifugation at 11500 g, 4°C, for 10 min and the supernatant was collected and passed through a PTFE 0.2 µm pore size syringe filter.

Quantification of carotenoids was performed by HPLC using a Shimadzu Prominence HPLC system (Shimadzu, Kyoto, Japan) consisting of two LC-20AD pumps, a DGU-20A degasser, a SIL-20AC automatic sample injector, CTO-20AC column thermostat and a Nexera X2 SPD-M30A photodiode-array detector. Chromatographic separations were carried out on a Phenomenex Synergi Hydro-RP 250 x 4.6 mm column with a particle size of 4 µm and a pore size of 80 Å. 20 μl aliquots of acetonic extract was injected to the column and the pigments were eluted by a linear gradient from solvent A (acetonitrile, water, triethylamine, in a ratio of 9:1:0.01) to solvent B (ethylacetate). The gradient from solvent A to solvent B was run from 0 to 25 min at a flow rate of 1 ml/min. The column temperature was set to 25 °C. Eluates were monitored in a wavelength range of 260 nm to 750 nm at a sampling frequency of 1.5625 Hz. Pigments were identified according to their retention time and absorption spectrum and quantified by integrated chromatographic peak area recorded at the wavelength of maximum absorbance for each kind of pigments using the corresponding molar decadic absorption coefficient (Jeffrey et al., 1997). The de-epoxidation index of the xanthophyll cycle components was calculated as (zeaxanthin + antheraxanthim)/(violaxanthin + anteraxanthin + zeaxanthin).

### Chemical treatments

For Asc supplementation, 1 mM Na-Asc (Roth GmbH) was added to the cultures, and measurements were carried out after a 2 h incubation period in the light.

For H_2_O_2_ treatments, the cell density was adjusted to 3 million cells/ml and 1.5 mM H_2_O_2_ (Sigma Aldrich) was added. The presented measurements were carried out 7 h following the addition of H_2_O_2_.

Catalase (5 µg/ml, from bovine liver, Sigma Aldrich) was added after a 30-min dark adaptation and the measurements were carried out after an additional 2 h incubation period in the dark with shaking.

### Western blot analysis

Protein isolation and western blot analysis were performed as in Vidal-Meireles et al., (2017). Specific polyclonal antibodies (produced in rabbits) against PsaA, PsbA, RbcL, LHCSR3, CP43, and PetB were purchased from Agrisera AB. Specific polyclonal antibody (produced in rabbits) against PSBO was purchased from AntiProt.

### NPQ measurements

Chlorophyll *a* fluorescence was measured using a Dual-PAM-100 instrument (Heinz Walz GmbH, Germany). *C. reinhardtii* cultures were dark-adapted for 30 min and then liquid culture containing 30 µg Chl(a+b)/ml was filtered onto Whatman glass microfibre filters (GF/B) that was placed in between two microscopy cover slips with a spacer to allow for gas exchange. For NPQ induction, light adaptation consisted of 30 min illumination at 530 µmole photons m^−2^ s^−1^, followed by 12 min of dark adaptation interrupted with saturating pulses of 3000 µmole photons m^−2^ s^−1^.

### Low-temperature fluorescence emission spectra (77K) measurements

Algal cultures containing 2 µg Chl(a+b)/ml were collected at several time points during the NPQ induction protocol. Subsequently, the sample was filtered onto a Whatman glass microfibre filter (GF/C), placed in a sample holder and immediately frozen in liquid N_2_. Low-temperature (77K) fluorescence emission spectra were measured using a spectrofluorometer (Fluorolog-3/Jobin–Yvon–Spex Instrument S.A., Inc.) equipped with a home-made liquid nitrogen cryostat. The fluorescence emission spectra between 650 and 750 nm were recorded with an interval of 0.5 nm, using an excitation wavelength of 436 nm and excitation and emission slits of 5 and 2 nm, respectively. The final spectra were corrected for the photomultiplier’s spectral sensitivity.

### Statistics

The presented data are based on at least three independent experiments. When applicable, averages and standard errors (±SE) were calculated. Statistical significance was determined using one-way ANOVA followed by Dunnett multiple comparison post-tests (GraphPad Prism 7.04; GraphPad Software, USA). Changes were considered statistically significant at p < 0.05.

## Acknowledgements

The authors thank Prof. Dr. Ralph Bock (MPI-MP Potsdam, Germany) and Dr. Petar Lambrev (BRC Szeged, Hungary) for discussions, Dr. Anikó Galambos (BRC Szeged, Hungary) for the assistance with ordering the mutants from the CLiP library, and for Dr. László Szabados (BRC Szeged, Hungary) for the possibility to use their CCD camera.

No conflicts of interest declared.

## Supplemental figures

**Supplemental Fig. S1.** Confirmation of the location of the CIB1 casette in the insertional CLiP mutant of *C. reinhardtii* (*LMJ.RY0402.058624*, named *Crvtc2-1*), affected in the *VTC2* gene. A, Physical map of the *VTC2* gene (obtained from Phytozome v12.1.6) with the CIB1 cassette insertion site in the *Crvtc2-1* mutant. Exons are shown in black, intron in light grey. Insertion site of the CIB1 cassette is indicated by triangle and the sequencing region is marked by arrows; B, Sequencing results using the Multiple Sequence Alignment by CLUSTALW (https://www.genome.jp/tools-bin/clustalw).

**Supplemental Fig. S2.** Characterization of the CC-4533 (wild-type), *Crvtc2-1* mutant, and *Crvtc2-1+VTC2* complemented *C. reinhardtii* lines. A, Cell volume. B, Chl(a+b) per million cells. C, Chl *a*/*b* ratio. D, Increase in Chl(a+b) concentration during culture growth in mixotrophic conditions (TAP) at moderate and high light (100 and 530 µmole photons m^−2^ s^−1^, respectively) and in photoautotrophic conditions (HSM) at 530 µmole photons m^−2^ s^−1^. E, Cellular Asc content after 24 h of growth under conditions corresponding to panel D. Data was analyzed by one-way ANOVA followed by Dunnett post-test: ××× p<0.05; ×× p<0.01; ××× p<0.001; ×××× p<0.0001 compared to the CC-4533 strain at the respective time-point and treatment. µE stands for µmole photons m^−2^ s^−1^.

**Supplemental Fig. S3.** Western blot analysis for the semi-quantitative determination of PsbA, CP43, PSBO, PsaA, LHCSR3, PetB and RbcL contents in *C. reinhardtii* cultures grown under photomixotrophic conditions at moderate light (100 µmole photons m^−2^ s^−1^) and photoautotrophic conditions at moderate and high light (100 and 530 µmole photons m^−2^ s^−1^ respectively). Samples of 1 µg Chl(a+b) were loaded, and the first four lanes (25, 50, 100, and 200% of the CC-4553 strain grown under photomixotrophic conditions at 100 µmole photons m^−2^ s^−1^) are for the approximate quantitation of the proteins. The graphics represent the densitometry analysis based on three independent experiments. Data was analyzed by one-way ANOVA followed by Dunnett post-test: ×<0.05, ××<0.01, ×××<0.001 compared to the TAP-grown CC-4533 culture. µE stands for µmole photons m^−2^ s^−1^.

**Supplemental Fig. S4.** NPQ kinetics induced by strong red light (530 µmole photons m^−2^ s^−1^) in cultures grown either in photomixotrophic conditions in TAP medium at 100 µmole photons m^−2^ s^−1^ or in photoautotrophic conditions in HSM medium at 530 µmole photons m^−2^ s^−1^. A and B, NPQ kinetics of the CC-4533 (wild type), *Crvtc2-1* and *Crvtc2-1+VTC2* complemented lines; C and D, NPQ kinetics of the *VTC2-amiRNA* and empty vector (*EV2*) lines. Data was analyzed by one-way ANOVA followed by Dunnett post-test: #<0.05, ##<0.01 compared to the untreated CC-4533 culture at the respective time-point. µE stands for µmole photons m^−2^ s^−1^.

**Supplemental Fig. S5.** Carotenoid contents of the *Crvtc2-1* mutant and the wild type (CC-4533) during NPQ induction by strong red light (530 µmole photons m^−2^ s^−1^). The cultures were grown in photomixotrophic conditions in TAP medium at 100 µmole photons m^−2^ s^−1^. A, Violaxanthin; B, antheraxanthin; C, zeaxanthin; D, β-carotene; E, lutein concentrations. Data was analyzed by one-way ANOVA followed by Dunnett post-test: × p<0.05, ×× p<0.01, ×××× p<0.0001 compared to the dark-adapted CC-4533 strain.

**Supplemental Fig. S6.** NPQ induction in the *stt7-9* mutant of *C. reinhardtii* and in *cw15-412*, used as a control strain, grown in TAP medium at 100 µmole photons m^−2^ s^−1^. A, NPQ kinetics, induced by 530 µmole photons m^−2^ s^−1^ of red light and followed by a recovery phase; B, de-epoxidation index, determined in the growth light, after 30 min of dark adaptation, following illumination with strong red light, and after the recovery phase, as indicated in the scheme in panel A. Data was analyzed by one-way ANOVA followed by Dunnett post-test: #<0.05, ####<0.0001 compared to the *cw15-412* strain at the respective time-point; ×× p<0.01, ×××× p<0.0001 compared to the dark-adapted *cw15-412* strain. µE stands for µmole photons m-2 s^−1^.

**Supplemental Fig. S7.** Carotenoid contents of the *Crvtc2-1* mutant and the wild type (CC-4533) during NPQ induction upon strong red light (530 µmole photons m^−2^ s^−1^). The cultures were grown in photoautotrophic conditions in HSM medium at 530 µmole photons m^−2^ s^−1^. A, Violaxanthin; B, antheraxanthin; C, zeaxanthin; D, β-carotene; E, lutein concentrations. Data was analyzed by one-way ANOVA followed by Dunnett post-test: × p<0.05, ×× p<0.01, ××× p<0.001, ×××× p<0.0001 compared to the dark-adapted CC-4533 strain.

**Supplemental Fig. S8.** Acclimation to 530 µmole photons m^−2^ s^−1^ of red light followed by recovery in CC-4533 and *Crvtc2-1* cultures, grown photoautotrophically in HSM medium at 100 µmole photons m^−2^ s^−1^. A, NPQ kinetics; B, De-epoxidation index; C, F_V_/F_M_ parameter measured after dark adaptation and after strong red light illumination. D, 684 nm/ 710 nm ratio of the 77K fluorescence spectra. Samples were collected at the growth light of 100 µmole photons m^−2^ s^−1^, after 30 min of dark-adaptation, at the end of the 30 min strong red light illumination, and 12 min after the cessation of actinic illumination, as indicated by arrows in the scheme in panel A. E, The effect of 1.5 mM H_2_O_2_ on NPQ induction in the CC-4533 strain; F, the effect of 1.5 mM H_2_O_2_ on NPQ induction in the *Crvtc2-1* mutant; G, the effect of catalase on NPQ induction in the CC-4533 strain; H, the effect of catalase on NPQ induction in the *Crvtc2-1* mutant. Data was analyzed by one-way ANOVA followed by Dunnett post-test: × p<0.05, ×× p<0.01, ×××× p<0.0001 compared to the dark-adapted CC-4533 strain. µE stands for µmole photons m^−2^ s^−1^.

## Notes

*Funding:* This work was supported by the Lendület/Momentum Programme of the Hungarian Academy of Sciences (LP-2014/19), the National, Research and Development Office (NN 114524, GINOP-2.3.2-15-2016-00026) and the Alexander von Humboldt Foundation (grants to S Z T). A V-M received fellowships from the SEB / Company of Biologists Travel Fund, US DOE Travel Award and from the International Society of Photosynthesis Research (sponsored by Wiley on behalf of Plant, Cell and Environment).

